# Species Identification by Bayesian Fingerprinting: A Powerful Alternative to DNA Barcoding

**DOI:** 10.1101/041608

**Authors:** Ziheng Yang, Bruce Rannala

## Abstract

A number of methods have been developed to use genetic sequence data to identify and delineate species. Some methods are based on heuristics, such as DNA barcoding which is based on a sequence-distance threshold, while others use Bayesian model comparison under the multispecies coalescent model. Here we use mathematical analysis and computer simulation to demonstrate large differences in statistical performance of species identification between DNA barcoding and Bayesian inference under the multispecies coalescent model as implemented in the BPP program. We show that a fixed genetic-distance threshold as used in DNA barcoding is problematic for delimiting species, even if the threshold is “optimized”, because different species have different population sizes and different divergence times, and therefore display different amounts of intra-species versus inter-species variation. In contrast, BPP can reliably delimit species in such situations with only one locus and rarely supports a wrong assignment with high posterior probability. While under-sampling or rare specimens may pose problems for heuristic methods, BPP can delimit species with high power when multi-locus data are used, even if the species is represented by a single specimen. Finally we demonstrate that BPP may be powerful for delimiting cryptic species using specimens that are misidentified as a single species in the barcoding library.

## Introduction

DNA barcoding was initially proposed as a fast and inexpensive approach to species identification. With a reference library of sequences for a “universal locus” from species that are identified *a priori*, biological specimens can be identified through calculation of the genetic distance between the query and sequences in the library (Hebert et al., 2003). The universal locus is typically mitochondrial cytochrome oxidase 1 (CO1) or cytochrome b (cytb), as mtDNA is easier to type than nuclear DNA from highly processed and degraded tissue. One particularly successful application of DNA barcoding is in forensics or “DNA detective” work, where DNA evidence is used to track illegal trade of wildlife (Alacs et al., 2010). Similarly adult life stages of insects with easily identifiable morphological features can be used to construct the reference library, and then DNA barcodes from juvenile or incomplete specimens can be matched to the library.

DNA barcoding has also been used for species discovery or species delimitation. This typically relies on determining a genetic distance threshold or ‘barcoding gap’. The query specimen is identified and assigned to an existing species in the library if the shortest pairwise sequence distance from the query to the sequence library is smaller than a pre-specified threshold. If the smallest distance exceeds the threshold, there will be a non-identification, which indicates that the specimen is from a new species, not represented in the library. The threshold may be arbitrarily chosen. For example, the ‘10x rule’ (Hebert et al., 2004) specifies the interspecific divergence to be at least 10 times as large as the intraspecific diversity. Dowton (2014) used 4% of cox1 divergence. More sophisticated methods generate a threshold by taking a database with a known taxonomy and minimizing the false-positive errors (incorrectly identifying a specimen as a new species) and false-negative errors (incorrectly lumping a specimen into another species) (Meyer and Paulay, 2005). For example, Ratnasingham and Herbet (2013) calculated a threshold of 2.2%. Methods have also been developed to “optimize” thresholds in empirical databases, such as Spider (Brown et al., 2012) and ABGD (Automatic Barcode Gap Discovery, Puillandre et al., 2012). However, different species have different population sizes and divergence times. As a result, one expects considerable overlap between intraspecific variation and interspecific divergence (Meyer and Paulay, 2005), so that there may not be a “one-size-fits-all” threshold, whatever method is used to determine the threshold. In a case study examining DNA barcoding performance in a diverse group of marine gastropods, Meyer and (2005) found that use of one threshold to delineate all species was particularly problematic for closely related species in taxonomically understudied groups.

In contrast, the multispecies coalescent (MSC) model (Rannala and Yang, 2003) naturally accommodates different population sizes and species divergence times. In theory it should be possible to delimit species even in extreme cases where the within-species diversity for some species is higher than the between-species divergence between other species.

Another potential advantage of the MSC-based approach is its ability to delimit cryptic species (specimens in the database that are distinct species but recognised as one species). Although the potential clearly exists, the performance of bpp to delimit cryptic species in practical data analysis has not been carefully examined. Indeed Collins and Cruickshank (2014) suggested that “it is questionable whether such statistics would be reliable due to the sampling and parameter estimation problems associated with taxon rarity in species delimitation methods.” We demonstrate that BPP can delimit cryptic species with high accuracy even if the species are under-sampled.

There has been much recent discussion about the impact of species under-sampling, or the rarity of species, on species delimitation (Lim et al., 2012; Collins and Cruickshank, 2014). Rarity indeed appears to be very common. For example, 48.5% of species in the African beetles library examined by Ahrens et al. (2016) were singletons. Many authors have considered species rarity to be a major challenge for species delimitation. For example, Lim et al. (2012) claim that many newly developed methods either implicitly or explicitly require that all species are well sampled. They argue that delimitation techniques should be modified to accommodate the commonness of rarity. Their conclusions are based on the intuition that species delimitation requires information about the within-species diversity relative to the between-species divergence, and such information will be hard to obtain if some species are under-sampled or are singletons. Furthermore, heuristic methods of species delimitation have indeed been found to suffer from poor species sampling. For example, in a case study of southern African beetles using cox1 sequences from >500 specimens and ~100 species, Ahrens et al. (2016) demonstrated that limited sampling effort can compromise species delimitation by the Generalized Mixed Yule Coalescent (GMYC) method (Fujisawa and Barraclough, 2013). The difficulty appears to lie more with the high sensitivity of GMYC to variable population sizes than with high proportions of singletons per se; GMYC appears to have difficuly generating reliable estimates of intra- vs. interspecies evolutionary parameters when some species are under-sampled.

We note that species rarity is naturally accommodated by BPP. It is simply a matter of information content and power, and BPP can make reliable inferences using multi-locus data even if a species is represented by a single specimen (singleton). New species have often been described based on rare specimens using morphology and it therefore appears self-evident that genetic data should also contain enough information to infer the species status of a rare specimen.

In this paper, we conduct simple computer simulations to demonstrate the differences between an MSC-based method (BPP) and heuristic barcoding methods for species delimitation. First, we show that in realistic scenarios it is impossible to identify a single distance threshold (barcoding gap) that allows correct species delimitation for all species. However, BPP can identify all species in this case without difficulty. Second, we illustrate the use of BPP to detect cryptic species. Finally we study the problem of rare specimens or singletons, and demonstrate that with multi-locus data BPP can delimit singleton species with high precision and high accuracy.

## One barcode gap for identifying all species may not exist

We simulate sequence data using the species tree of figure 1a, with 1 + 10 sequences from *A*, 10 *B*s, 1 + 10 *C*s, 10 *Ds*, 10 *Es*, and 1 *F.* The 10 sequences each from species *A-E* (50 sequences in total) are used as the ‘library’, while the 1 *A*, 1 *C* and 1 *F* are used as three ‘query’ sequences (denoted *A*_1_, *C*_1_, and *F*). We generate one locus, of 1000bp, by simulating the gene trees under the MSC (Rannala and Yang, 2003) and evolving sequences along the gene tree. The program MCCOAL, which is part of the BPP package (Yang, 2015), was used for the simulations. The number of replicate simulated datasets was 100.

**Fig. 1.**
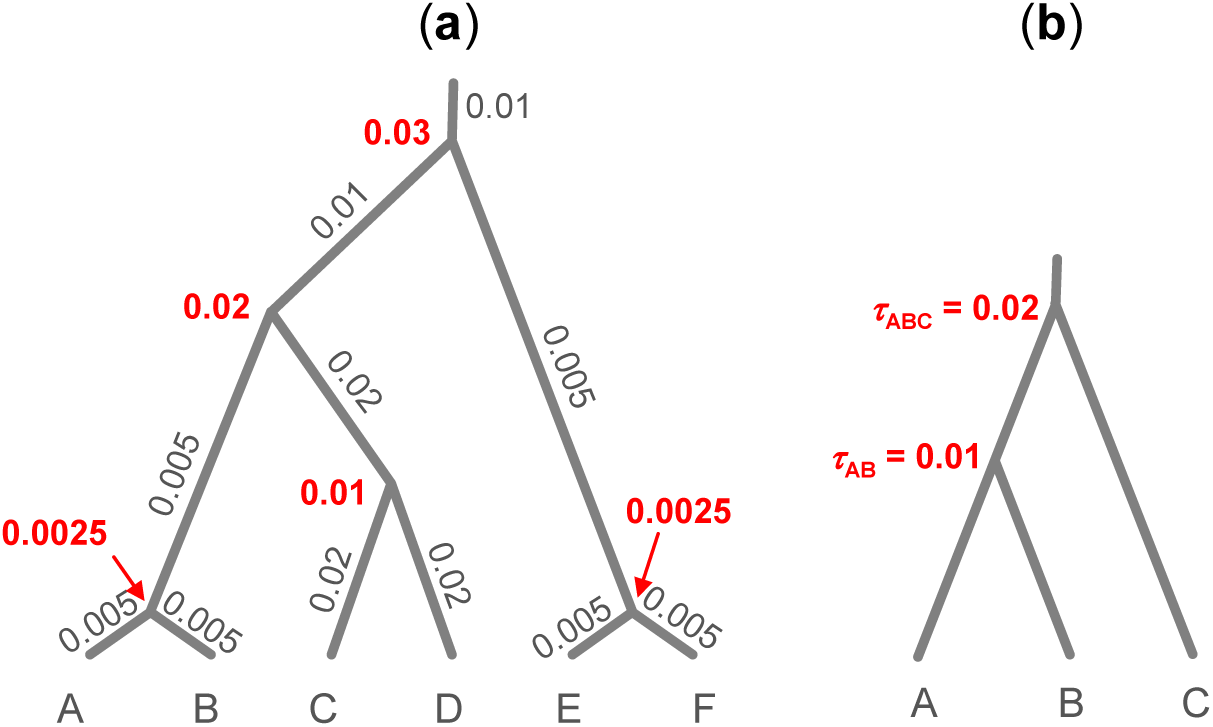
Species trees used in simulating sequence alignment under the MSC, with branches drawn using the species divergence times (τs). In (a), the species divergence-time parameters are shown in red bold font next to the internal nodes: τ_*AB*_ = τ_*EF*_ = 0.0025, τ_*CD*_ = 0.01, τ_*ABCD*_ = 0.02, and τ_*ABCDEF*_ = 0.03, while the population size parameters are shown along the branches: *θ*_*A*_ = *θ*_*B*_ = *θ*_*AB*_ = *θ*_*E*_ = *θ*_*F*_ = *θ*_*EF*_ = 0.005,*θ*_*C*_ = *θ*_*D*_ = *θ*_*CD*_ = 0.02, and *θ*_*ABCD*_ = *θ*_*ABCDEF*_ = 0.01. *In* (b), the parameters are τ_*AB*_ = 0.01, τ_*ABC*_ = 0.02, and *θ* = 0.01 for all populations. Both τs and *θ*s are measured by the expected number of mutations per site.

We analyzed the simulated data using both the DNA barcoding gap and the BPP program. Note that BPP will never separate one population into two species but may merge different populations into one species. The reversible-jump Markov chain Monte Carlo (rjMCMC) algorithm (Yang and Rannala, 2010) explores different delimitation models, which correspond to different groupings of populations into the species, as well as different phylogenetic relationships among the delimited species. The analysis assumes 8 populations: A, B, C, D, and E, and the three query sequences. We use Prior 3 for the species-tree models, which assigns uniform probability (1/8) for 1, 2, …, and 8 species (Yang, 2015). The priors on parameters are *θ ~* G(2, 200), with mean 0.01, and *τ*_ABC_ ~ G(3, 100), with mean 0.00333. The shape parameters 2 and 3 are relatively small, indicating that the gamma priors are fairly diffuse. The prior means are set to be equal to the true values.

For each simulated dataset (which consists of 53 sequences for one locus), sequences *A*_1_, *C*_1_ and *F* were queried against the sequence library made up of the remaining 50 sequences. Let *d(A*_1_, *A) = min{d(A*_1_, *A*_*i*_), *i* = 2, …, 10} be the smallest distance from the query *A*_1_ to species *A*, and define *d(C*_1_, *C*) and *d(F, E)* accordingly. *A*_1_ is correctly assigned to species *A* if *d(A*_1_, *A)* is smaller than the distance threshold, while *F* is correctly assigned to be a species distinct from *E* if *d(F, E)* is larger than the distance threshold. Thus to assign *A*_*1*_ and *C*_1_ correctly to species *A* and *C*, respectively, one would prefer a large distance threshold, and to assign *F* correctly into a distinct species from species *E*, one would like a small distance threshold. It will be impossible to assign all three queries correctly if *d(F, E)* ≤ *d(A*_1_, *A)* or if *d(F, E)* ≤ *d(C*_1_, *C).* This happened in 28 out of the 100 replicate datasets. For example in one dataset, *d(A*_1_, *A) = d(A*_1_, *A_2_)* = 0.001, *d(C*_1_, *C) = d(C*_1_, *C*_5_) = 0.004, and *d(F, E) = d(F, E_2_) =* 0.003. With the within-species distance (0.004 for *C*) being greater than the between-species distance (0.003 between *F* and *E)*, it is impossible to use one distance threshold to make correct assignments. Whatever the distance threshold, it will be impossible to avoid both the false positive error of claiming *A*_*1*_ or *C*_1_ as a new species (a non-identification) and the false negative error of lumping *F* into species *E*. In the other datasets, it is theoretically possible to choose a threshold that will allow correct assignment of all three queries, but mis-assignments may still occur if an imperfect threshold is chosen.

Fig. 2 shows the proportion (among 100 replicate datasets) of correct species assignments when each dataset is analyzed using a fixed distance threshold. For example, at the distance threshold of 0.003 (three differences per kb), *A*_*1*_ is correctly assigned to species *A* in 96% of datasets, *C*_1_ is correctly assigned to species *C* in 70% of datasets, and *F* is correctly assigned to a species distinct from species *E* in 39% of datasets.

**Fig. 2.**
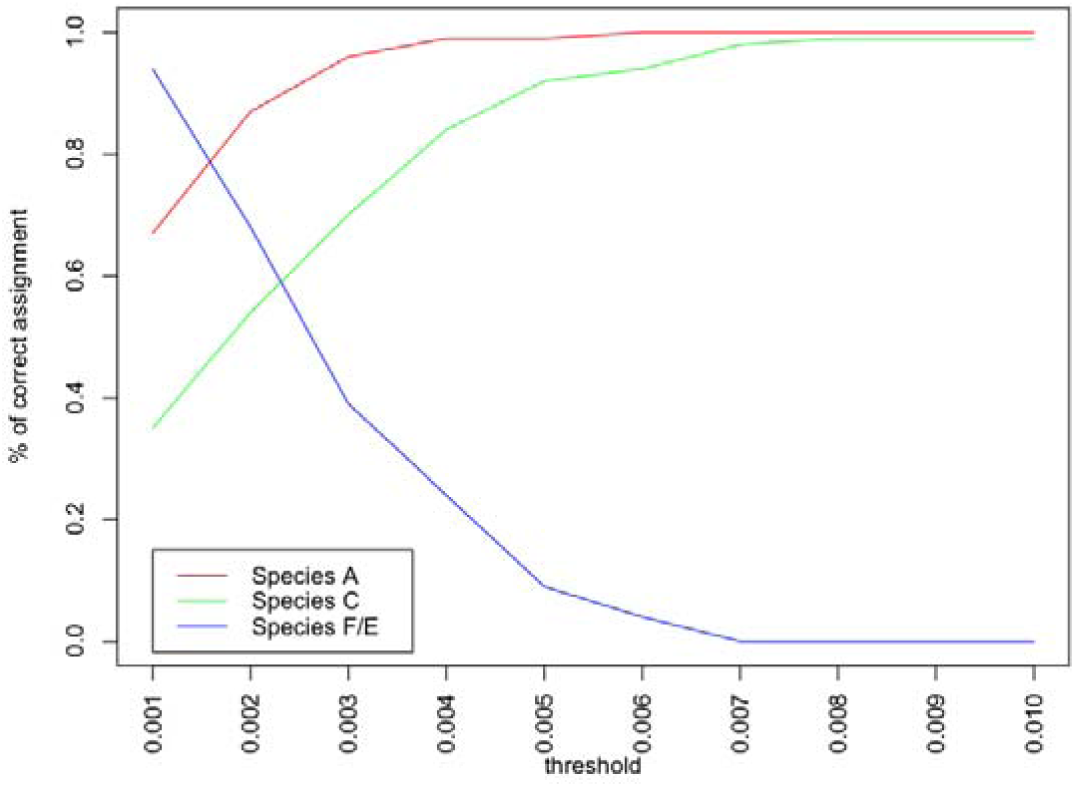
The proportion of correct species assignments when the distance threshold is fixed at different values, averages over 100 simulated replicate datasets.

The same datasets of one single locus were also analyzed using BPP and the resulting posterior probabilities of assignments are summarized in fig. 3 for data. Here P(*A*_1_ *A*) is the posterior probability that population *A*_1_ and *A* are (correctly) grouped into one species, to the exclusion of all other populations. The average posterior probability was 0.81 for correctly assigning *A*_1_ to species *A*, and was 0.17 for recognizing it as a new species (over-splitting). The average posterior probability was 0.71 for correctly assigning *C*_1_ to species *C*, with the false positive rate of oversplitting to be 0.29. The average posterior probability for correctly identifying *F* as a new species was 0.68, with the false negative rate of incorrectly lumping it with species *E* to be 0.31. Although the information is weak with a single locus, BPP is clearly outperforming the threshold method. High posterior probabilities (>0.9) for an incorrect assignment are very rare (less than 1% on average).

**Fig. 3.**
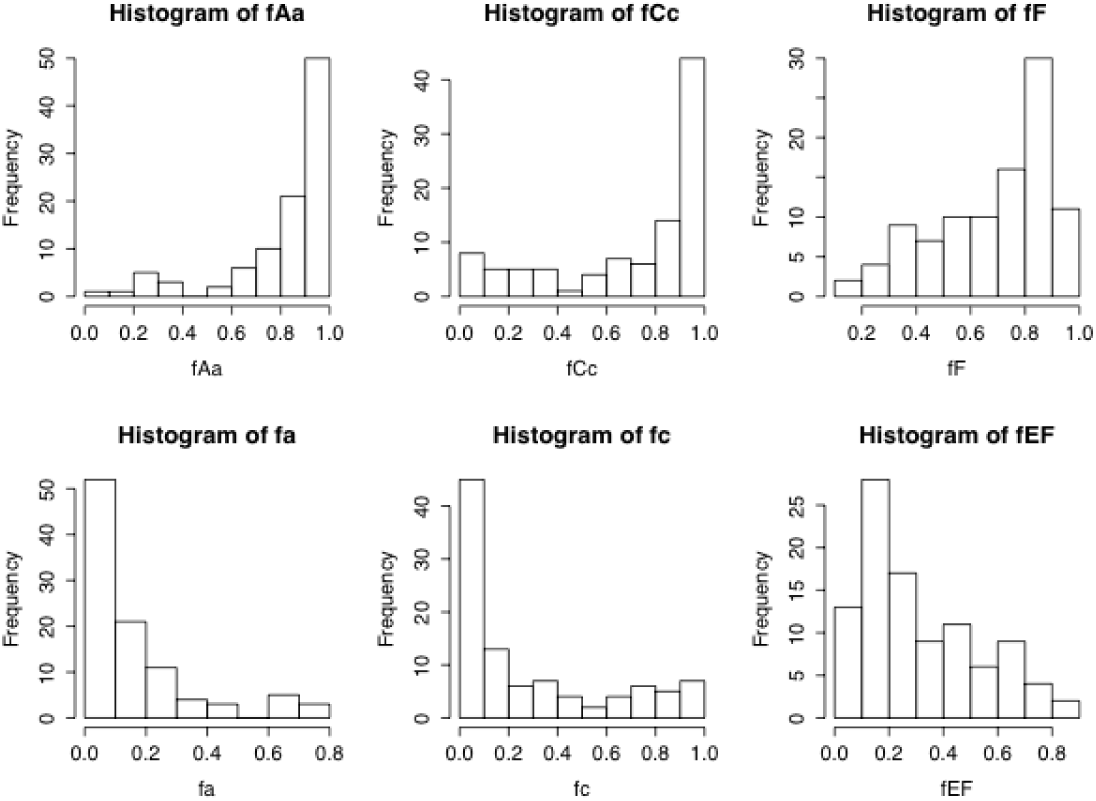
The histograms of posterior probabilities for correctly assigning *A*_1_*A*, *C*_1_*C*, and *F* into one species, and for incorrectly assigning *A*_1_, *C*_1_, *EF* into one species by BPP in datasets of one single locus.

Increasing the number of loci from 1 to 10 led to increased posterior probabilities for correct assignments and to reduced error rates. The posterior probabilities for correctly assigning *A*_1_ to species *A, C*_1_ to species *C*, and *F* to a distinct species from *E* are shifted towards 1 (fig. 4a-c), while the posterior probabilities for incorrectly assigning *A*_1_ or *C*_1_ to distinct species are shifted towards 0 (fig. 4d-f). Query *F* is identified as a distinct species with posterior probability 1.0, and the error rate for lumping *E* and *F* is 0 in every dataset.

**Fig. 4.**
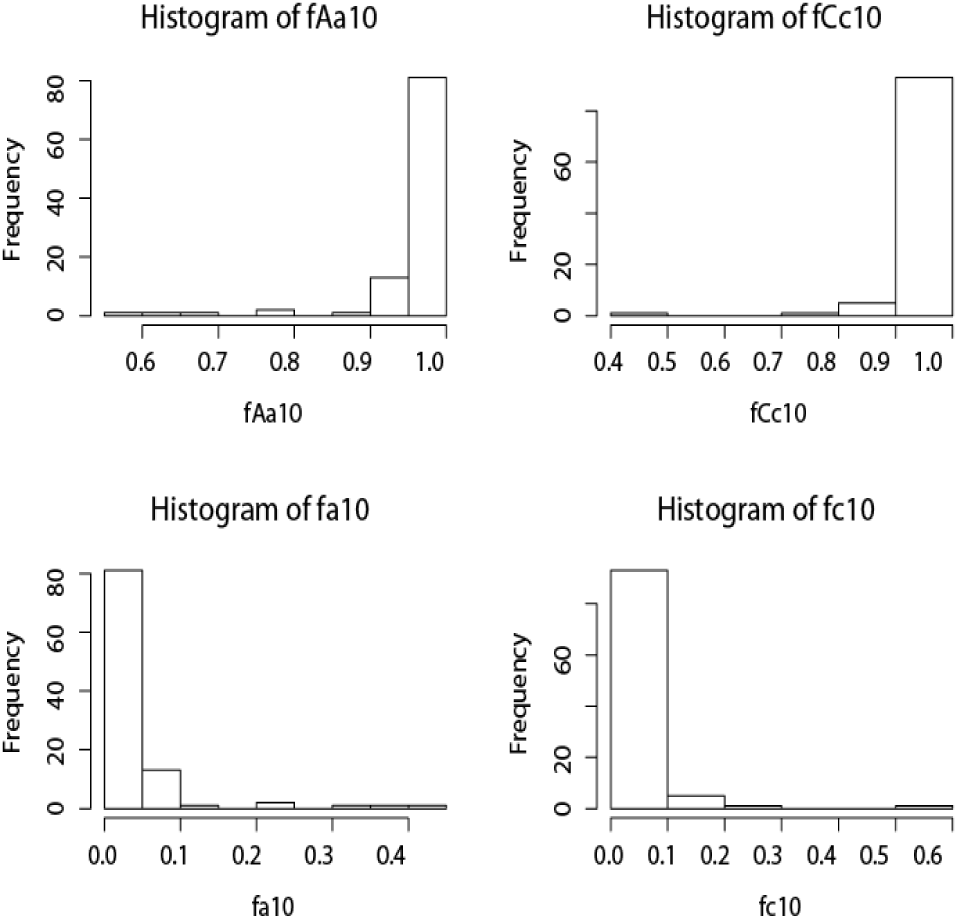
The histograms of posterior probabilities for species assignments by BPP in datasets of 10 loci. See legend for fig. 3. P(*F*) = 1 and P(*EF*) = 0 in every dataset so that those plots are not shown.

## Identifying cryptic species

To examine the performance of BPP in identifying cryptic species we simulated sequence data using the species tree for 3 species of figure 1b, with 4 sequences (two diploid individuals) from each species. The parameter values were *θ* = 0.01 for all populations, *τ*_*AB*_ = 0.01 and *τ*_*ABC*_ = 0.02. A and B represent distinct cryptic species, misidentified as one species in the library. Each locus was 1000bp. The number of loci was either 2 or 10. The number of replicate simulations was 100. The BPP analysis assumed 5 populations *(A*_1_, *A_2_, B*_1_, *B*_2_, *C*), with each individual from A and B treated as a separate population. We assign Prior 3 for the species-tree models, with uniform probability (1/5) for 1, 2, …, 5 species (Yang, 2015). The priors on parameters are *θ* ~ G(1, 100) and τ_ABC_ ~ G(4, 200). These are diffuse priors with the means equal to the true values.

The true model in this case has 3 species, with the phylogeny *((A, B*), *C*), and with *A*_1_ and *A*_*2*_ grouped into one species *(A)*, and *B*_1_ and *B*_2_ into another *(B).* With 2 loci, 82% of the simulated datasets produced a maximum *a posteriori* (MAP) model with 3 species that matched the true model. The histogram for the posterior probability for the correct model (which separates *A* and *B* into distinct species and also infers the correct species phylogeny) is shown in fig. 5a. While this probability is >70% in most datasets, it is low in many other datasets, reflecting the low information content in data of one single locus. With 10 loci, 97% of the simulated datasets produced a MAP model with 3 species that matched the true model. The histogram for the posterior probability for the correct model is shown in fig. 5b. The shift towards 1 relative to that of fig. 5a reflects the dramatic increase in the information content in data of 10 loci.

**Fig. 5.**
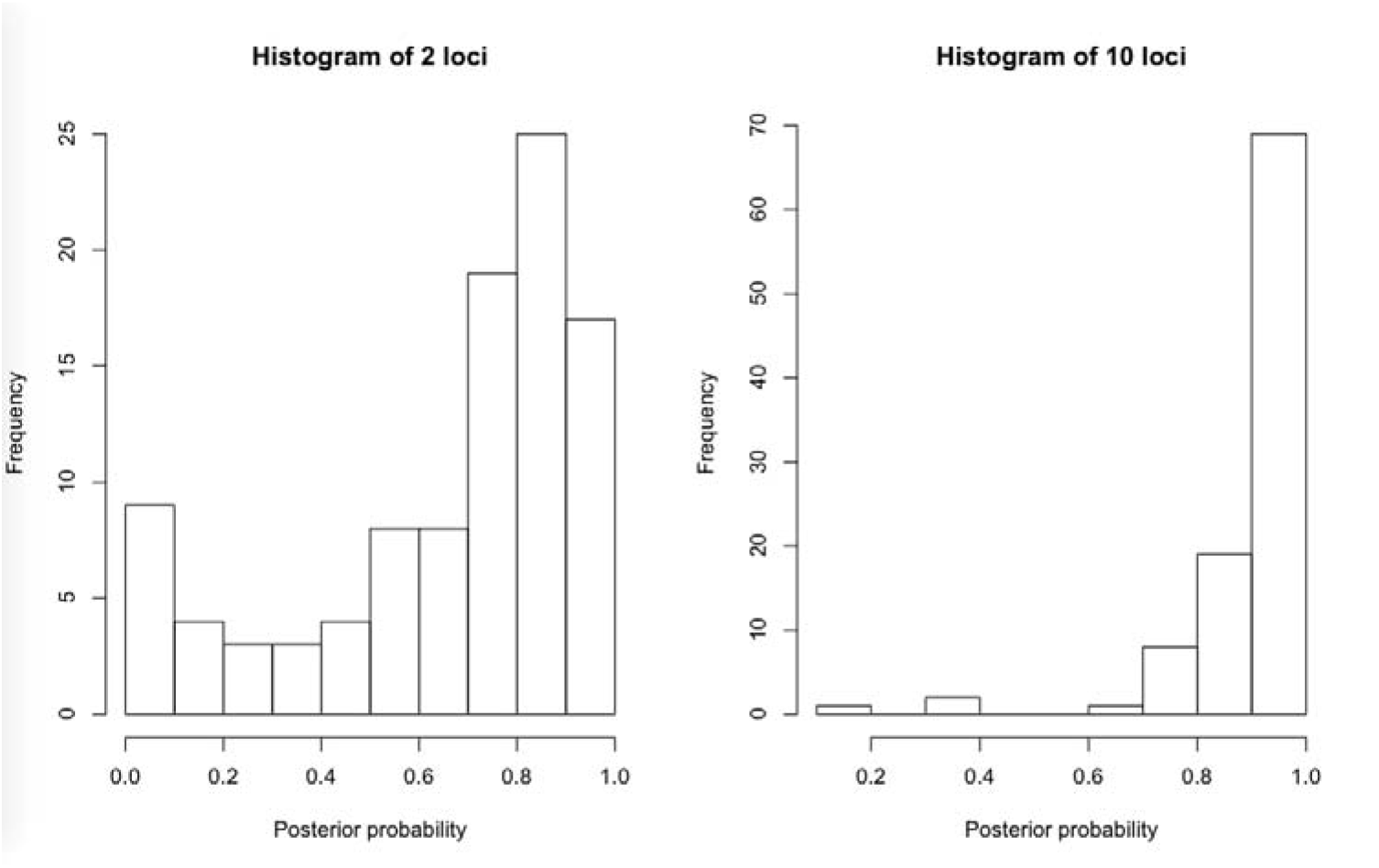
The histograms of posterior probabilities for correctly inferring the cryptic species status as well as the species phylogeny by BPP using datasets of 2 and 10 loci. There are 3 species in the dataset, with *A* and *B* representing two cryptic species.

If a posterior probability of 0.90 is used as a cut-off to choose a model, the error rate is 1% for 2 loci and 0% for 10 loci. With a 0.90 posterior probability cut-off the power to identify the true model is 17% with 2 loci and 69% with 10 loci. The average posterior probability Pr(*A*_1_ *A*_2_) for grouping *A*_1_ and *A*_2_ into one species was 0.83 with 2 loci and 0.94 with 10 loci, and the average posterior probability Pr(*B*_1_*B*_2_) for grouping *B*_1_ and *B*_2_ into one species was 0.85 with 2 loci and 0.94 with 10 loci. The distributions of posterior probabilities of the *A*_1_ *A*_2_ and *B*_1_*B*_2_ groupings with either 2 or 10 loci are shown in fig. 6. Additional loci may be needed to infer the true model with almost complete certainty.

**Fig. 6.**
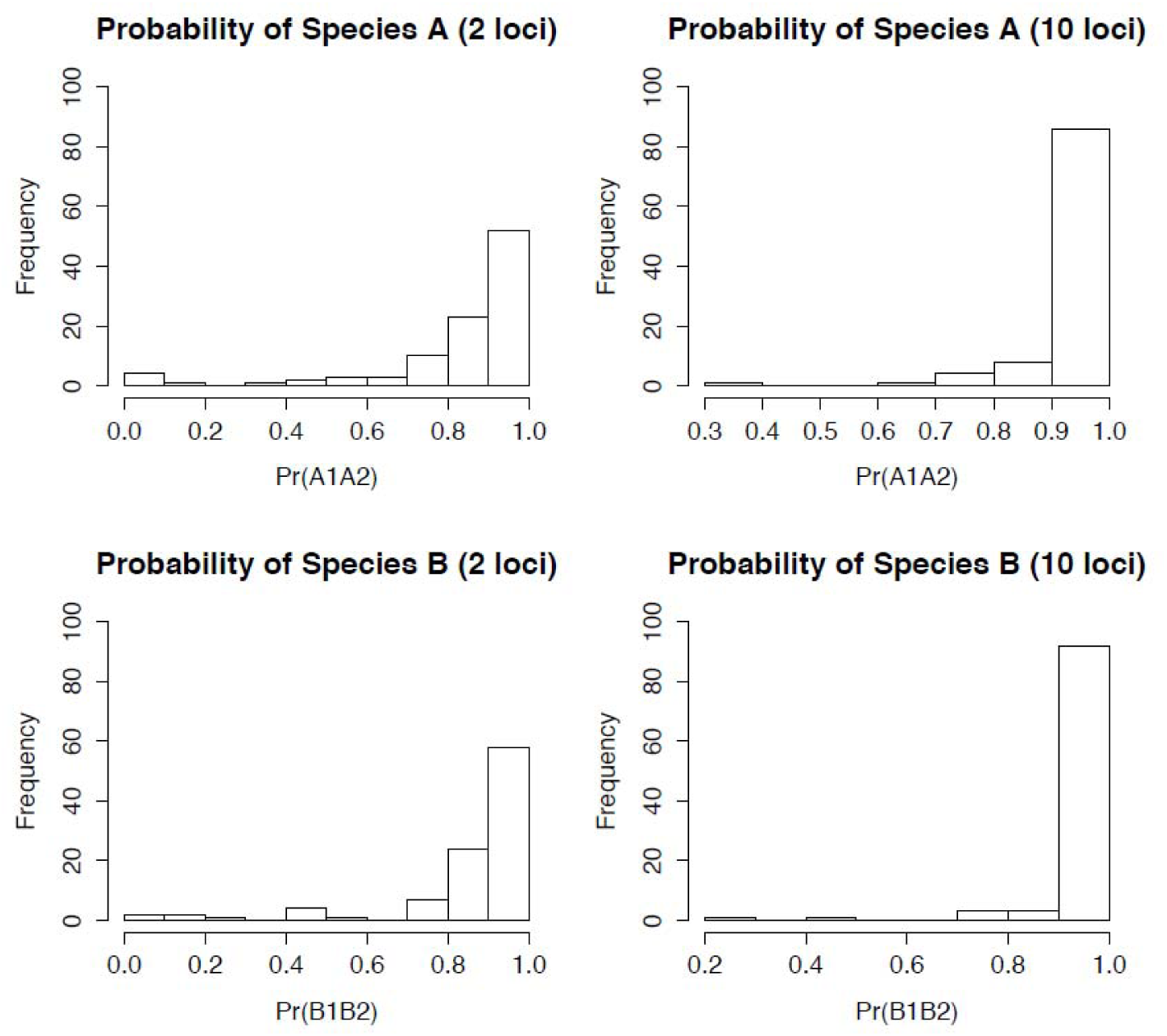
The histograms of posterior probabilities for correctly identifying cryptic species by BPP using datasets of 2 and 10 loci.

## Identifying rare species

To examine the performance of BPP in identifying rare species we simulated data on the species tree of figure 1b, with 1 *A*, 10 *B*s, and 10 *C*s. The one *A* sequence represents one specimen (a singleton) from a haploid species. We are interested in whether BPP can correctly infer *A* to be a distinct species when sequence data from multiple loci are available. Note that having two *A* sequences (as would be available if the species is diploid) will make the task easier. The parameter values are *θ* = 0.01 for all populations, *τ*_*AB*_ = 0.01 and *τ*_*ABC*_ = 0.02. Each locus was 1000bp. The number of loci was either 2 or 10. The number of replicates was 100. The BPP analysis assumes 3 populations *(A, B, C).* We assign Prior 3 for the species-tree models, which assigns uniform probability (1/3) for 1, 2, and 3 species (Yang, 2015). The priors on parameters are *θ ~ G(1, 100) and τ*_*ABC*_ ~ G(4, 200).

The true model in this case is 3 species, with the species tree ((*A, B*), *C*). The MAP model was the true model for all simulated datasets with either 2 loci or 10 loci. In all datasets, three species were delimited with posterior probability greater than 0.95, whether 2 or 10 loci are analyzed. The average posterior probability of 3 species was 0.998 with 2 loci and 1.000 with 10 loci. The power of inference is very high in this case compared with the simulation of cryptic species, because multiple individuals (10 sequences) are available from species *B*.

## Discussion

We suggest that species identification and discovery using genetic sequence data be best viewed as a statistical inference problem, given the stochastic nature of the coalescent and the process of sequence evolution. Species assignment by BPP under the MSC makes an efficient use of information in the sequence distances, or in the greater between-species divergence than within-species polymorphism. However, BPP in addition uses information in the gene trees and branch lengths, even though the gene trees at each individual locus may involve considerable uncertainties and sampling errors. The method accommodates the fact that some species have large population sizes and thus show greater within-species diversity, and that some species diverged recently so that the between-species divergence may not necessarily exceed the within-species diversity. By using a formal modeling framework and incorporating information about contemporary and ancestral population sizes available from multilocus sequence data one avoids the need to specify subjective distance thresholds as in DNA barcoding. Another advantage of the statistical modeling methods such as BPP over heuristic methods such as DNA barcoding is that they provide measures of uncertainties in the form of posterior probabilities.

Lim et al. (2012; see also Collins and Cruickshank, 2014) claim that “in studies using coalescence much of the evidence for species limits comes from coalescence points, which are by definition lacking for rare species…” This intuition is faulty, as can be seen from our simulation results showing that BPP identified the singleton species with higher power, even though the single sequence from the singleton species cannot provide any within-species coalescent points. Indeed, the simulation of Zhang et al. (2011, fig. 3) showed that BPP can assign species correctly with 10 or 50 loci (depending on the mutation rate or sequence divergence level) even if a single sequence is sampled from each population at every locus so that estimation of intra-species diversity is not possible for either species. Lim et al. (2012) went on to recommend several approaches for identifying statistical ‘outliers’ for use in species recognition. Those suggestions are neither valid nor relevant.

Dowton et al. (2014) suggested that coalescent-based species delimitation methods can be used to make more accurate specimen identifications than single-locus DNA barcoding. However, Collins and Cruickshank (2014) suggest that “to benchmark the efficiency and accuracy of species delimitation methodologies, it should now be a priority to highlight exemplar data sets—empirical and/or simulated—for which MSC methods clearly outperform simpler mtDNA analyses.” In this study we have generated several such exemplar cases by simulation and showed that simple distance threshold does not work well.

Collins and Cruickshank (2014) further discussed the computational burden of species delimitation methods based on the MSC. While algorithmic improvements are being made (Rannala and Yang, 2016), it may be interesting to note that BPP recovered the true model (species delimitation and species phylogeny) with near certainty with a moderate number of loci such as 10 or 20. Thus there is probably no need to use the whole genome to infer species status. Inference methods such as BPP can be practical and powerful for smaller numbers of loci and individuals. Similarly there may not be a need to analyze 500 species, say, in one combined analysis, to delimit species. It may be reasonable to analyze divergent groups of species as separate datasets, thus further reducing the computational burden involved. Collins and Cruickshank (2014) stated that they “agree that researchers should be looking into ‘smarter’ methods for taxon identification, but only in cases where the ‘dumb’ methods have first been comprehensively shown not to work.” Here we show that for the species tree that we considered it was impossible for the ‘dumb’ barcoding method to make correct identifications for all species in about 30% of datasets. The BPP method, on the other hand, produced reasonable results with one locus, favouring the correct assignment and never supporting a wrong assignment with high posterior probability. With 10 loci, it had both high precision and high accuracy. We suggest that the ‘dumb’ method failed dramatically in this case. We suggest that “comprehensively” showing that the method does not work is unnecessary since a practical ‘smart’ method is already available that does not have the pathological behaviour and will achieve accurate inferences with sufficient numbers of loci. We note that even the two dumbest methods one could imagine, one assigning every query specimen to a new species, and the other assigning every query specimen to the most common species in the library, cannot be shown “comprehensively” to fail, because they will provide perfect answers in a substantial proportion of datasets.

## Acknowledgments.

This study is supported by a grant from Biotechnological and Biological Sciences Research Council (BBSRC, UK) to Z.Y.

